# Circulating U13 small nucleolar RNA as a candidate biomarker for Huntington’s disease

**DOI:** 10.1101/2022.04.22.489178

**Authors:** Silvia Romano, Carmela Romano, Martina Peconi, Alessia Fiore, Gianmarco Bellucci, Emanuele Morena, Fernanda Troili, Virginia Cipollini, Viviana Annibali, Simona Giglio, Rosella Mechelli, Michela Ferraldeschi, Liana Veneziano, Elide Mantuano, Gabriele Sani, Andrea Vecchione, Renato Umeton, Franco Giubilei, Marco Salvetti, Rosa Maria Corbo, Daniela Scarabino, Giovanni Ristori

**Affiliations:** Department of Neurosciences, Mental Health and Sensory Organs, Sapienza University of Rome, SantAndrea Hospital, Rome, Italy; Department of Human Neurosciences, Sapienza University of Rome, Rome, Italy; Department of Biology and Biotechnology, Sapienza University, Rome, Italy; Department of Experimental Medicine, Sapienza University, Policlinico Umberto I, Rome, Italy; San Raffaele Roma Open University and IRCCS San Raffaele, Rome, Italy; Ospedale San Giovanni Battista, ACISMOM. Rome, Italy; CNR Institute of Translational Pharmacology, Rome, Italy; Department of Psychiatry, Fondazione Policlinico Universitario Agostino Gemelli IRCCS, Rome, Italy; Department of Neuroscience, Section of Psychiatry, Catholic University of the Sacred Heart, Rome, Italy; Surgical Pathology Units, Department of Clinical and Molecular Medicine, SantAndrea Hospital, Sapienza University, Rome, Italy; Department of Informatics and Analytics, Dana-Farber Cancer Institute, Boston, MA, USA; Massachusetts Institute of Technology, Cambridge, MA, USA; Harvard School of Public Health, Boston, MA, USA; Weill Cornell Medicine, New York City, NY, USA; IRCCS Istituto Neurologico Mediterraneo (INM) Neuromed, Pozzilli, Italy; Department of Biology and Biotechnology, La Sapienza University, Rome, Italy; CNR Institute of Molecular Biology and Pathology, Rome, Italy; Neuroimmunology Unit, IRCCS Fondazione Santa Lucia, Rome, Italy

**Keywords:** [14] All Clinical Neurology, [91] All Genetics, [164] Huntington’s disease

## Abstract

**Background and Objectives:** Fluid biomarkers are a recent field of interest in Huntington disease (HD). We focused on small circulating RNAs from plasma of subjects with prodromal (pre-HD) and overt disease by a two-stage approach: an unbiased investigation by an array method and a validation study to quantify a significant small nucleolar RNA.

**Methods:** Through Affymetrix Gene-Chip-miRNA-Array we performed an exploratory study on 9 HD patients, 8 healthy subjects (HS) and 5 psychiatric patients (PP; who share drugs with HD patients, to control for iatrogenic effects). Through real time PCR we validated the results in an independent population of 24 HD patients, 15 pre-HD, 24 PP, 28 Alzheimer’s disease (AD) patients (added to control the disease-specificity of our finding) and 23 HS. A bioinformatic analysis was also performed to interpret our finding.

**Results:** The microarray results showed a significant signal for U13 small nucleolar RNA (SNORD13) that was increased in plasma of HD patients compared to controls (fold change, 1.54, p =0.003 HD vs. HS, and fold change 1.44 p = 0.0026 HD vs. PP). In the validation population the significant increase in HD patients was evident compared to both pre-HD and the three control groups (p<0.00001). The plasma levels of SNORD13 correlated with the status of mutant huntingtin carrier and the disease duration (respectively R=0.69; p<0.000001; R=0.49; p=0.015). Through receiver operating characteristic (ROC) curve analysis, we showed high accuracy of plasmatic SNORD13 in discriminating HD patients from pre-HD and control groups (AUC=0.963), outperforming values reported in another study for intrathecal or plasmatic mutant huntingtin and neurofilament light chain as biomarkers of overt HD. The bioinformatic analysis on SNORD13 interactome and pathway analysis showed enrichments for factors involved in nuclear functions beyond the ribosome biogenesis.

**Discussion:** We report the unprecedented finding of a potential role of small nucleolar RNAs in HD. Circulating SNORD13 seems a good biomarker for clinical purposes. It seems to be specific for HD and to peripherally report a plausible ‘tipping point’ in the pathogenic cascade at neuronal level, possibly paving the way for new therapeutic targets.

## INTRODUCTION

Huntington disease (HD) is an inherited neurodegenerative disease due to a CAG trinucleotide repeat expansion in the first exon of the HTT gene, encoding the huntingtin protein. HD is a progressive, incurable disease, with typical adult onset, related to CAG repeat length, characterized by motor impairment, cognitive dysfunctions and psychiatric symptoms. Disease-modifying treatments for HD are in development and the identification of easily measurable biomarkers is crucial to predict progression, to monitor the effects of novel drugs and to obtain cues on the pathogenic cascade at neuronal level.

Fluid biomarkers are quantified in body fluids, ideally with minimal invasiveness, good accuracy, and high discriminatory power. Cerebrospinal fluid (CSF) has been a focus of interest, as a proxy of Central Nervous System (CNS) pathophysiology: recent studies identified reliable biomarkers such as CSF mutant Huntingtin (mHTT), used as an outcome measure for therapeutic approaches ^1^, and neurofilament light chain (NfL)^2^, which showed very good predictive power for disease manifestation ^3,4^. Biomarkers based on complex techniques or CSF may be of limited use because of their invasiveness, high costs, or need of specialized personnel.

A more recent focus was given to more easily measurable biomarkers, coming from peripheral leucocytes and plasma, that are cheaper, less invasive and potentially more adept to obtain longitudinal profiles. Among these, the leukocyte telomere length (LTL) shorten remarkably in pre-HD, being a possible measure of time to clinical onset ^5^. Changes in histone variant pγ-H2AX, as a component of the DNA damage responses, in peripheral blood mononuclear cells (PBMCs), proved to be an informative, potentially reversible biomarker in pre-HD ^6^. In other studies, PBMCs from HD patients were used to study mHTT and the gene expression profiles as predictors of disease progression ^7,8^. Plasma NfL resulted reliable in monitoring disease progression, though less sensitive than CSF NfL ^4^. Other studies reported informative results on plasma levels of oxidative stress markers, metabolic markers and immune system products ^9^. Recent studies on small non-coding RNAs in plasma from HD patients led to several investigations on circulating micro-RNAs (miRNA) ^10–12^. Our recent work found that hsa-miR-323b-3p was upregulated in individuals with mHTT mutation ^13^. In this context, we obtained results on small nucleolar RNAs (snoRNA), that were not previously studied in HD; we considered both snoRNAs predictive significance as biomarkers and their possible role in disease pathogenesis.

SnoRNAs are a class of non-coding small guide RNAs, most of which direct the chemical modifications of other RNA substrates, including ribosomal RNAs (rRNAs) and spliceosomal RNAs. Moreover, some snoRNAs are involved in the regulation of alternative splicing and post-transcriptional modification of mRNA^14^. The Homo sapiens U13 snoRNA ([SNORD13](https://www.ncbi.nlm.nih.gov/gene?cmd=Retrieve&dopt=Graphics&list_uids=692084)), 104 nucleotides long, is member of the Box C/D family of small nucleolar ribonucleoprotein; it can form base-pair interactions with the 3’ portion of 18S rRNA, and is involved in the processing of this rRNA^15^. Our study is aimed at reporting the changes of the circulating SNORD13 in HD as an informative biomarker for disease management and a possible relay of central pathogenic events.

## MATERIALS AND METHODS

### Study Population

Participants in the study were enrolled at the Center for Experimental Neurological Therapies, Unit of Neurology (patients with positive test for HD, patients with probable Alzheimer’s disease and healthy subjects) and at the School of Medicine and Psychology, Unit of Psychiatry (patients with psychiatric disorders), S. Andrea Hospital, Department of Neurosciences, Mental Health, and Sensory Organs, Sapienza University of Rome, Italy. The study was approved by the local Ethics Committee, and a written consent was obtained from all participants, according to the principles expressed in the Declaration of Helsinki. The operators were unaware of the disease state of each sample during processing and statistical analysis. Eligible subjects for this study were patients with a positive genetic test for HD, healthy subjects, patients with diagnosis of schizophrenia or bipolar disorders treated with antipsychotic drugs, and patients with diagnosis of probable Alzheimer’s disease. Exclusion criteria were pregnancy, breastfeeding, and severe systemic illnesses or conditions. Based on preliminary results, we estimated that groups of 12 cases and 12 control subjects would have been sufficient to detect a variation of at least ±40% for the parameter, assuming a standard deviation (SD) of 34% with a power of 90%, using a two sided significance level of 0.05.

### Plasma Preparation and Affymetrix Gene Chip miRNA Array

Blood samples were obtained by venous punctures in ethylenediaminetetraacetic acid (EDTA) tubes for plasma preparation. Plasma was obtained by centrifugation (1.500 g for 15min at 4°C), a few 500 μl aliquots of supernatant was stored at −80°C. RNA was extracted using Plasma/Serum Circulating RNA Purification Kit (NORGEN) following the manufacturer’s instructions. RNA quality and purity were assessed with the use of the RNA 6000 Nano Assay on Agilent 2100 Bioanalyzer (Agilent). Briefly, 500 ng of total RNA was labeled using FlashTag Biotin HSR (Genisphere) and hybridized to GeneChipR miRNA 2.0 Arrays. The arrays were stained in the Fluidics Station 450 and then scanned on the GeneChip R Scanner 3000 (Affymetrix,). The statistical analysis was performed by Transcriptome Analysis Console (TAC) software (Thermo Fisher Scientific). In order to survey the presence of outliers that could impact the dataset, a principal component analysis (implemented in R) was performed to identify possible outliers that needed to be excluded. MicroRNA probe outliers were defined per manufacturer’s instructions (Affymetrix), and further analysis included data summarization, normalization, and quality control using the web-based miRNA QC Tool software (Affymetrix). The raw dataset is available in the Gene Expression Omnibus (GEO) repository (GSE167630).

### Real Time PCR analysis

RNAs extraction was performed using miRNeasy Serum/Plasma kit according to the manufacturer’s protocol (Qiagen). U13 snoRNA analysis was performed by quantitative reverse transcription-PCR (qRTPCR). cDNA was synthesized with TaqMan™ MicroRNA Reverse Transcription Kit (Life Technologies) according to the manufacturer’s instructions, using a U13 snoRNA-specific primer (manufacturer provided). Quantitative Real Time PCR was performed an ABI 7300 Real-time PCR System (using custom TaqMan® Small RNA Assay (Life Technologies), according to the manufacturer’s instructions. Relative quantitation of U13 snoRNA was calculated by the delta Ct method using U6 snoRNA as endogenous control ^16^. Replicate assays of same sample were completed to calculate the inter-assay variation. The average standard deviation (SD) as calculated by measuring plasma SNORD13 levels of a sample repeated over three different assays was 0.035%. Thus, assuming a normal distribution, samples differing in average SNORD13 levels by as little as 0.069% (1.96 X SD) should be distinguishable by this method at the 95% confidence interval ^17^.

### Statistical Analysis

Statistical analyses were performed using Partek Genomic Suite software (miRNA Array data), GraphPad Prism v9 and R (The R Project for Statistical Computing) v3.6.3. Data normality was assessed through Shapiro-Wilk test. Nonparametric tests were used to compare the distribution of SNORD13 plasma levels between patients and controls. Non-parametric tests or linear regression were used to evaluate the distribution of the SNORD13 plasma levels across age, sex, and CAG repeat number. Spearman’s correlation was computed to assess the linear relationship between SNORD13 plasma levels and disease duration or UHDRS-TMS. Data for variables of interest were available for all the subjects. Significance was taken at p<0.05 and ROC curve analysis was performed through the R easy ROC web interface. Optimal cut-point was identified according to the Youden Index method. Sample size estimation settings were as follows: Type I error: 0.05; Power: 0.8; Allocation ratio=1; AUC was derived from the HD/pre-HD ROC curve.

### Interactome construction and analysis

The molecular interactions of SNORD13 were searched in three databases: (1) the RNA interactome database (RNAinter v4.0) ^18^ at http://www.rna-society.org/rnainter/, which includes more than 41 million predicted or experimentally validated interactions; (2) SNOdb, the largest repository on snoRNAs biological annotation and manually curated snoRNA-RNA interactions ^19^; (3) RNACT, a database of genome wide predicted protein–RNAs interactions ^20^. Non-human interactors were excluded. Protein-protein interactions were derived from querying STRING database ^21^ for high-confidence interactions (score>0.7) of SNORD13-interacting proteins. The global SNORD13 interactome was mapped through Cytoscape ^22^. Pathway enrichment analysis was performed through the STRING app in Cytoscape. Graphical plotting was performed through the ggplot2 package in R ^23^.

## RESULTS

We performed an exploratory microarray study of whole noncoding RNA expression profiles in plasma from 9 patients with Huntington’s disease (HD, mean age of 48.25 ± 10.47) and 13 controls, including 8 healthy subjects (HS, mean age of 49.17 ± 11.79) and 5 psychiatric patients (PP, mean age of 50.25±11.47), with schizophrenia or bipolar disorder. The HD patients were treated with antipsychotic drugs, the most effective treatment option also in PP (we included the PP control group to minimize a possible iatrogenic impact of neuroleptic drugs on the profile of the small non-coding RNAs). The microarray results indicate that SNORD13 level was increased in plasma of HD patients compared to both control groups (fold change, 1.54, p =0.0003 HD vs. HS, and fold change 1.44 p = 0.0026 HD vs. PP; Figure 1A).

**Fig. 1:**
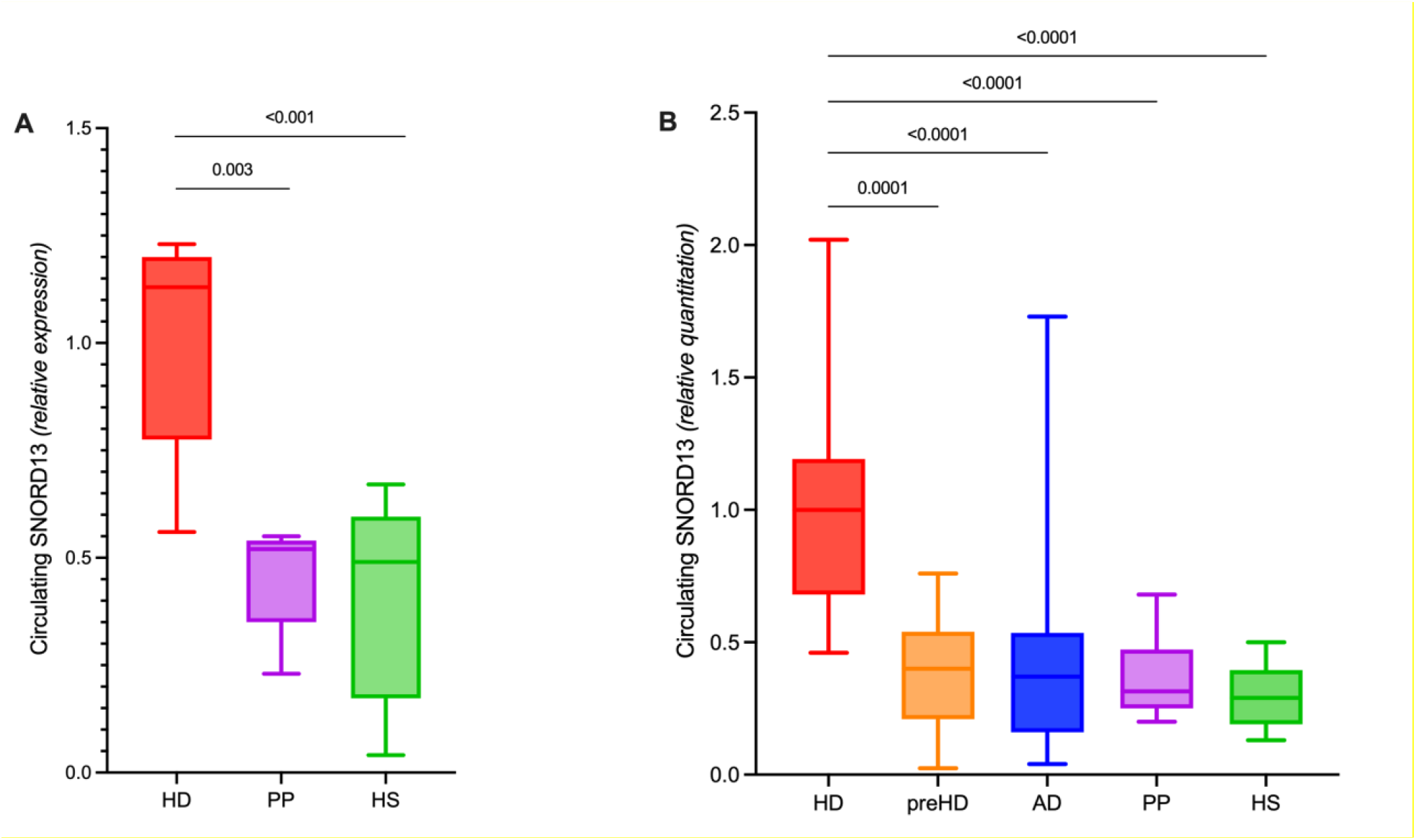
HD-specific signature of circulating SNORD13. (A) Relative expression of circulating SNORD13 measured through microarray in a cohort of HD patients, psychiatric patients (PP) and healthy subjects *(Mann-Whitney test)*. (B) Relative quantitation of circulating SNORD13 measured trough RT-PCR in an independent cohort of HD patients, premanifest mHTT carriers (pre-HD), people with Alzheimer’s Disease (AD), psychiatric patients (PP) and healthy subjects (HS) (*Kruskal-Wallis test)*.

To validate this result, SNORD13 plasma levels were quantified by Real Time PCR in five cohorts of subjects: 23 Healthy Subjects (HS), 24 symptomatic HD patients (HD), 15 pre-manifest HD (pre-HD), 24 Psychiatric Patients (PP) and 29 Alzheimer’s disease patients (AD). The last group was added to control the disease-specificity of our finding. Demographic and clinical characteristics of each group are presented in Table 1. No significant relationship was observed between SNORD13 plasma levels and age/sex at blood sampling in any group, except for an excess of female in HS (p=0.04). No relationship was observed between SNORD13 levels and CAG repeat length in pre-HD or HD subjects (Table 2). Our analysis showed a statistically significant increase in the plasma level of SNORD13 in HD patients, that clearly segregates patients with overt disease (HD) from controls and pre-HD subjects. The changes in the plasma level of SNORD13 in HD patients was highly significant (p<0.00001) compared to both pre-HD and the 3 control groups (HS, PP and AD; Fig. 1B, Table 2).

**Table 1:** Demographic and clinical characteristics of mHTT carriers and controls.

|  | Healthy subjects<br>n=23 | Pre-manifest<br>mHTT<br>carriers<br>n=15 | HD patients<br>n=24 | Psychiatric<br>patients<br>n=24 | AD<br>patients<br>n=29 |
| --- | --- | --- | --- | --- | --- |
| Age (years) | 50.3 ± 13.4<br>(range 26-76) | 36.4 ± 12.4<br>(range 21-59) | 53.6±11.5<br>(range 28-75) | 48.1±13.1<br>(range 20-67) | 67.6±5.3<br>(range 56-76) |
| Sex (males, %) | 40.9 | 26.6 | 69.6 | 66.7 | 63 |
| CAG<br>Median (range) | / | 42.8 (40-49) | 42.8 (40-47) | / | / |
| UHDRS-TMS | / | 1.1±2.1 | 30.2±16.0 | / | / |
| HD duration | / | / | 6.2±3.7 years<br>(range 1-14) | / | / |

**Table 2.** **Circulating SNORD13 in HD and control groups** (referring to Figure 1B).

| Group (n) | SNORD13 relative<br>quantitation<br>(median, 1st and 3rd<br>quartiles) | p-value<br>vs HS | Covariate analysis |  |  |
| --- | --- | --- | --- | --- | --- |
|  |  |  | SNORD13/Age<br>(p-value) | SNORD13/Sex<br>(p-value) | SNORD13/CAG<br>number (p-value) |
| HS (23) | 0.29 (0.19-0.41) | / | 0.21 | 0.04 |  |
| pre-HD (15) | 0.40 (0.21-0.54) | 0.1 | 0.48 | 0.44 | 0.37 |
| HD (24) | 0.96 (0.66-1.14) | <0.0001 | 0.53 | 0.33 | 0.77 |
| PP (24) | 0.32 (0.25-0.47) | 0.24 | 0.99 | 0.9 | / |
| AD (29) | 0.37 (0.16-0.56) | 0.43 | 0.19 | 0.84 | / |

A positive linear correlation was observed between circulating SNORD13 and disease duration in HD patients (r=0.49, p=0.015) (Fig. 2A). A significant relationship was also present between plasmatic SNORD13 and the UHDRS clinical score in mHTT carriers (r = 0.69, p <0.0001, Fig. 2B), while it was not found to be significant considering only overt HD patients (not shown).

**Figure 2:**
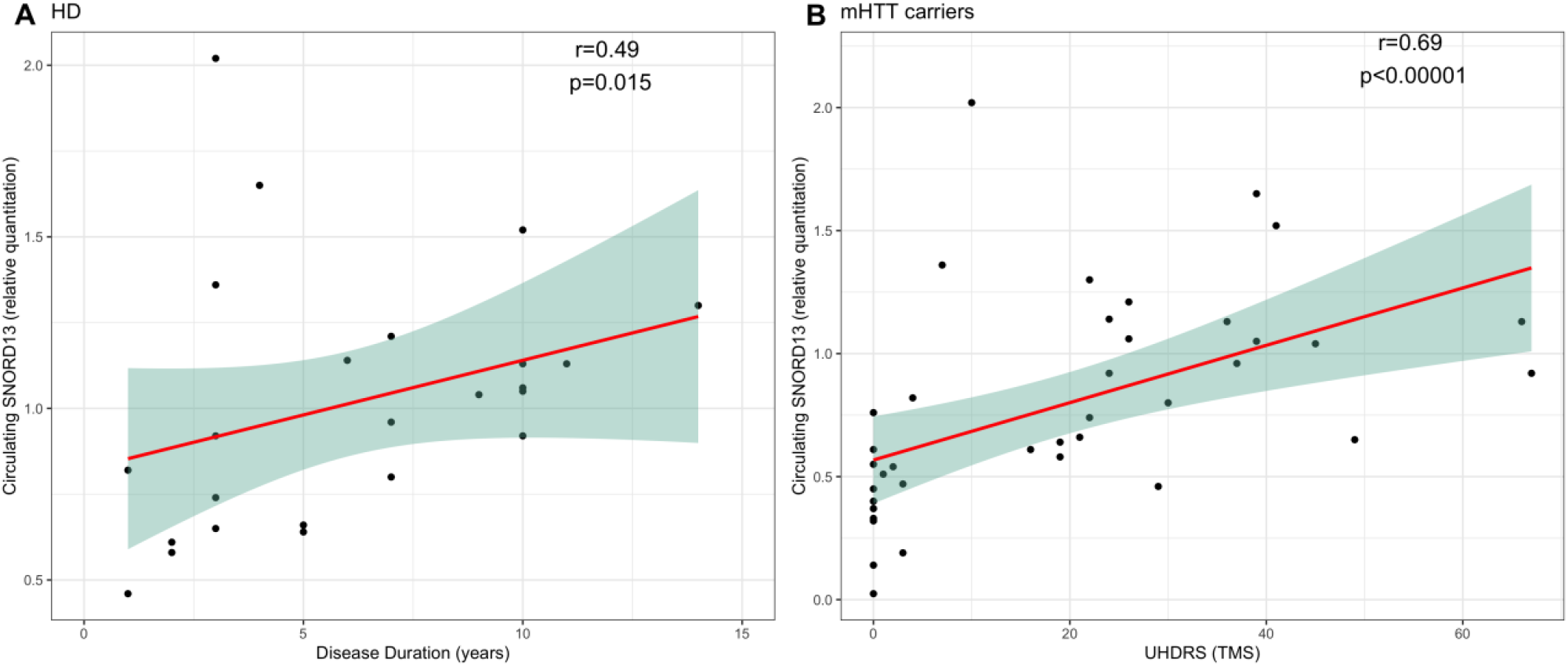
**Correlation between SNORD13 plasma levels and (A) disease duration in HD patients, and (B) Unified Huntington’s Disease Rating Scale Total Motor Score (UHDRS- TMS) in mHTT carriers** *(Spearman’s correlation)*.

Next, we assessed the accuracy of plasmatic SNORD13 as a biomarker of overt HD through receiver operating characteristic (ROC) curve analysis. In discriminating symptomatic HD patients from pre-symptomatic HTT mutation carriers, SNORD13 displayed extremely high accuracy (AUC=0.963 [0.911-1.000], figure 3A); setting the cut-point of SNORD13 levels at 0.58 allows to identify HD with 95.88% sensitivity and 86.7% specificity. We estimated a sample size of 3 HD and 3 pre-HD to validate SNORD13 validity as a biomarker in the clinical setting. Moreover, SNORD13 appeared to be of potential utility in distinguishing manifest HD patients from control groups (AD, PP, HS; AUC=0.953 [0.914-0.992]; figure 3B), also including pre-HD among controls (AUC=0.955 [0.918-0.991]; Figure 3C) and, to a lesser extent, in identifying HTT mutation carriers (AUC: 0.811 [0.719-0.904]; Figure 3D). We also compared the diagnostic performance of SNORD13 with the one of mHTT and NfL from peripheral blood and CSF, as described in Byrne et al., 2018 ^2^. While CSF mHTT level is the gold-standard diagnostic test to identify HTT mutation carriers (as expected), the plasmatic level of SNORD13 seems to outperform both mHTT and plasmatic or intrathecal NfL as a biomarker of clinically overt HD in our findings (Table 3).

**Table 3.** Diagnostic ability of SNORD13 compared with NFL and mHTT performance from the work by Byrne et al., 2018 (AUC derived from ROC curve analysis).

|  | Controls vs mHTT carriers | pre-HD vs HD |
| --- | --- | --- |
| CSF mHTT | 1.000 | 0.778 |
| CSF NFL | <u>0.933</u> | 0.914 |
| Plasma NFL | 0.914 | 0.931 |
| Plasma SNORD13 | 0.811 | <u>0.963</u> |

**Figure 3:**
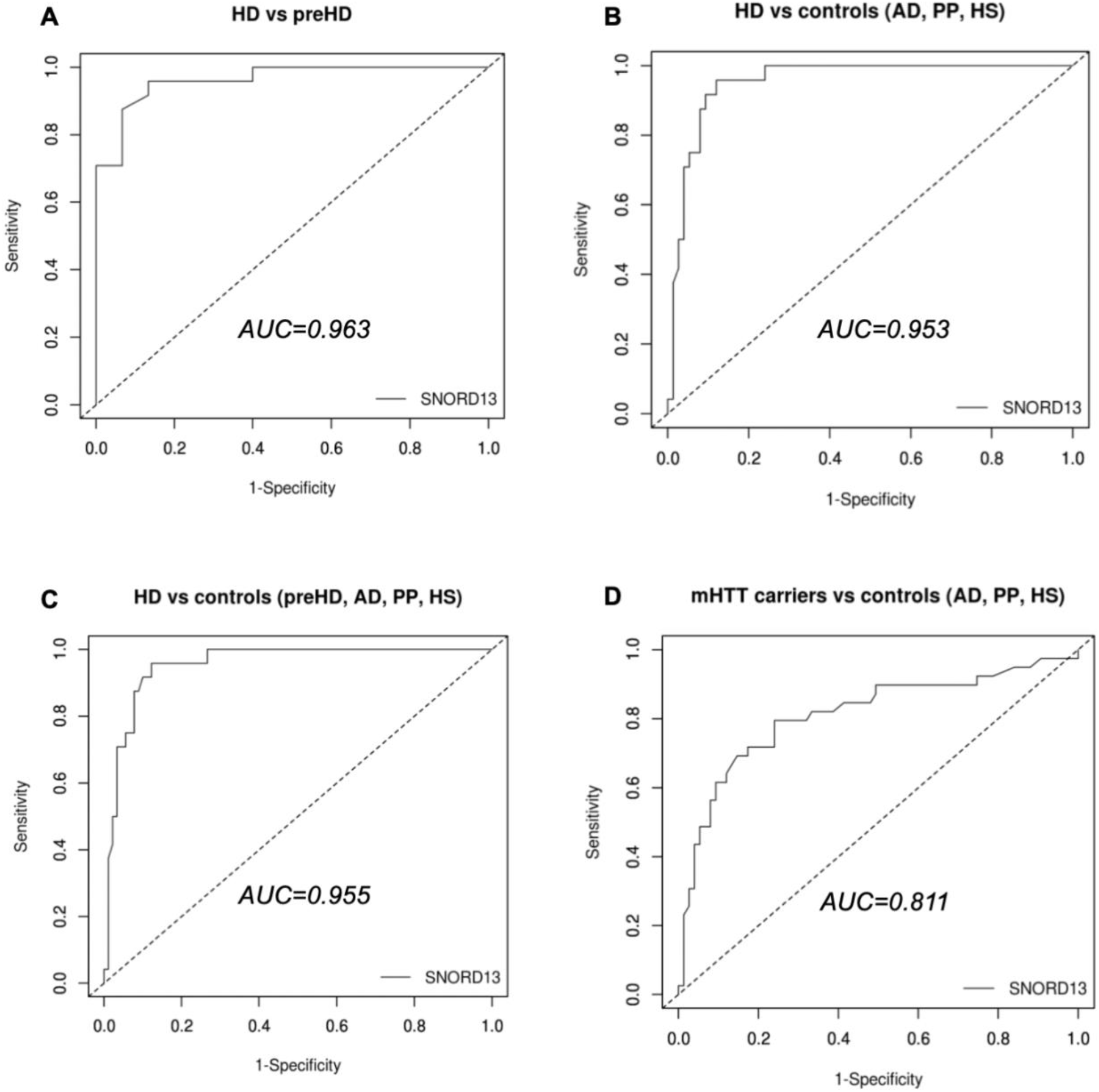
Assessment of SNORD13 as a biomarker of HD. ROC curves for discrimination between (A) symptomatic HD and premanifestomatic HD (pre-HD); (B,C) symptomatic HD and control groups; (D) mHTT carriers (HD and pre-HD) and control groups.

Finally, to investigate the biological landscape of action of SNORD13, we constructed its interactome. We retrieved U13 snoRNA interactions with proteins (including known RNA-binding proteins, RBPs, and Transcription Factors, TF), snoRNAs, miRNAs and rRNAs by interrogating the databases RNAinter ^19^, SNOdb ^20^ and RNAct ^21^ and additionally, we mapped the intra-network protein-protein interactions through STRING ^22^. The final network was formed by 456 SNORD3-interacting nodes: 258 TFs, 86 known RBPs, 91 proteins, 1 lncRNA, 3 miRNAs, 13 other snoRNAs and the 18s Rrna ribosomal subunit (Figure 4A and supplementary table 1). Pathway analysis revealed an enrichment of processes involved in transcriptional regulation and RNA metabolism (figure 4 B-D), referred to molecules mostly located in the nucleus and involved in genomic organization (Figure 4E). Of interest for HD is the emergence of NGF-stimulated transcription as associated with SNORD13 activity, suggesting a direct implication in neurodegenerative processes, as well as the interaction with molecules involved in the DNA damage response that was already implicated in disease pathogenesis and in a peripheral biomarker ^6,7,24^.

**Figure 4:**
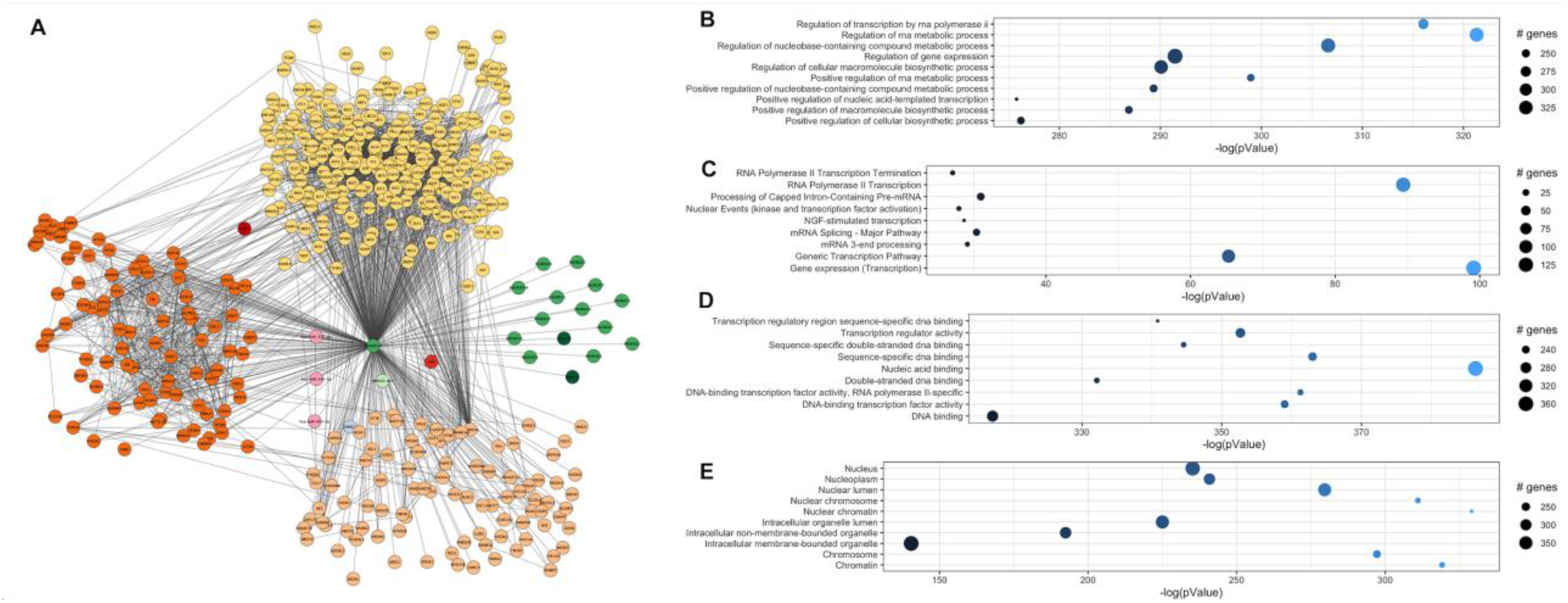
SNORD13 interactome and pathway analysis. (A) SNORD13 interaction map. (B-E) Top 10 results of enrichment analysis of SNORD13 interactors in: (B) Gene Ontology Biological Processes; (C) Reactome pathways; (D) Gene Ontology Molecular Function; (E) Gene Ontology Cellular Compartment.

## DISCUSSION

Our study highlights the unprecedented finding of a potential role of small nucleolar RNAs in HD. The main points related to this result are the following: (i) a circulating SNORD13 is significantly increased in the overt disease compared to the prodromal phase of HD; (ii) the levels of this snoRNA are comparable between healthy individuals and pre-HD status; (iii) this finding seems specific for HD, since three groups of control (healthy people, PP sharing drugs with HD patients, and AD patients) showed normal, comparable values; (iv) the plasma levels of SNORD13 seems to efficiently mark the natural history of HD, correlating with the status of mHTT carrier and the disease duration; (v) the above points (iii) and (iv) suggest that increased levels of SNORD13 in blood mirror pathogenic events in the CNS (though this fact awaits formal demonstration), possibly paving the way for new therapeutic targets; (vi) SNORD13 may become a good peripheral biomarker for clinical purposes in HD; it is indeed easily measurable, inexpensive to test, possibly linked to the pathophysiology of the disease, and reliably quantifiable (being consistent, accurate, sensitive, specific and reproducible) ^9^. A possible hypothesis to explain our data is that the elevated plasma level of SNORD13 in HD patients may be due to the nucleolar stress caused by the presence of mutant RNAs that carry an expanded CAG repeat (expanded CAG RNAs). In recent years, increasing attention has been directed toward the understanding of pathogenic mechanisms due to mHTT: several studies support the hypothesis that expanded CAG RNAs induce apoptosis by activating the nucleolar stress pathway in both HD patients and transgenic models of disease ^25,26^. Specifically, it has been shown that expanded CAG RNA competes with nucleolin (a multifunctional protein that is mainly localized in the nucleolus and is involved in various steps of ribosome biogenesis) for rRNA promoter, leading to reduction of rRNA expression, nucleolar stress and apoptosis via p53 and activation of the downstream signaling cascade, including mitochondrial cytochrome c release and caspase activation ^27–30^.

However, considering the SNORD13 interactome, we cannot rule out the possibility that other pathogenic mechanisms might be related to changes in the small nucleolar RNA. Indeed, beyond ribosomal biogenesis and RNA metabolism, SNORD13 appears to be involved in a wide range of genomic activities associated with HD pathophysiology (Figure 4, Supplementary Table 1): chromatin remodeling via histone acetylation and methylation (*SIRT6, EP300, TAF1, CHD1, KDM5A-B etc*.); telomere length maintenance (*DKC1,HRNPU, VPRBP)* ^5^; DNA repair and damage response (*ERCC6, BAZ1B, FANCD2, TOPBP1)* ^6,24^; direct modulation of Homeobox (*HOXA1-7, DLX1-2, OTX2* and others) and Zinc-finger proteins (*SNAI2, SP1-2, ZMIZ2* and others) ^31^. We also identified three miRNAs (hsa-miR-455-5p, hsa-miR-342-3p, hsa-miR-377-3p) that may cooperate with U13 snoRNA in regulating gene expression, as being possibly affected in HD; stochastic computational analyses might help further explain these relationships, as piloted in Multiple Sclerosis pathogenesis modeling ^32^, in COVID epidemic waves stability evaluation^33^, and in many other complex problems.

Irrespective of the mechanism(s) possibly linking mHTT to SNORD13 and the source of this small nucleolar RNA, we can speculate that the rising of the plasma level of SNORD13 in HD patients may peripherally report a ‘tipping point’ in the pathogenic cascade at neuronal level, while normal levels may mark a ‘molecular pre-manifest status’ in diseases evolution. In fact, when we used published data ^2^ on CSF mHTT and plasma or CSF NfL as benchmarks, to compare the performance of plasma SNORD13 as biomarker, we found that it outperforms both CSF mHTT and plasmatic or intrathecal NfL as reporter of overt HD, supporting its potential value as a peripheral cue of central pathogenic processes.

This study does not allow to directly benchmark SNORD13 against circulating NFL or CSF mHTT because we did not assay them in this pilot study. The other limitations of the present work fall within those of a pilot study: the sample size was informative, although future studies in larger groups may help to demonstrate the generalizability of the findings; moreover, longitudinal data showing the dynamics of circulating SNORD13 over time are missing, and future studies with repeated assays in the same subjects and an adequate number of pre-manifest and manifest mHTT carriers have been planned.

In summary, circulating SNORD13 could be a clinically actionable asset in future trials as a peripheral reporter of a ‘functional reserve’ at neuronal level, as well as a hint for new therapeutic targets. In general, peripheral small RNAs may offer potential advantages in terms of disease-specificity compared to other approach, such as circulating NFL, that was already reported as a promising peripheral biomarker for many neurological diseases ^3^. Future personalized therapies for different phases of HD course and possible trials with etiologic approaches in pre-manifest subjects may pose novel needs, that will plausibly be fulfilled by circulating biomarkers, with SNORD13 among the promising components of the neurogeneticist’s toolkit.

## Acknowledgements

We would like to thank the patients and their families and the healthy controls who took part in our study. This study was funded by a research grant from Sapienza University of Rome.

## REFERENCES

1. Tabrizi S. J., Leavitt B. R., Landwehrmeyer G. B. et al. Targeting Huntingtin Expression in Patients with Huntington’s Disease. N Engl J Med 2019, 380, 2307–2316. doi: 10.1056/NEJMoa190090.

2. Byrne L. M., Rodrigues F.B., Blennow K. et al. Neurofilament light protein in blood as a potential biomarker of neurodegeneration in Huntington’s disease: a retrospective cohort analysis. Lancet Neurol 2017;16, 601–609. doi: 10.1016/S1474-4422(17)30124-2.

3. Gaetani L., Parnetti L., Calabresi P. & Di Filippo M. Tracing Neurological Diseases in the Presymptomatic Phase: Insights from Neurofilament Light Chain. Front. Neurosci. 2021; 15:549. doi: 10.3389/fnins.2021.672954.

4. Scahill R. I., Zeun P., Osborne-Crowley K. et al. Biological and clinical characteristics of gene carriers far from predicted onset in the Huntington’s disease Young Adult Study (HD-YAS): a cross-sectional analysis. Lancet Neurol 2020; 19, 502–512. doi: 10.1016/S1474-4422(20)30143-5.

5. Scarabino D., Veneziano L., Peconi M., Frontali M., Mantuano E., Corbo R.M., Leukocyte telomere shortening in Huntington’s disease. J. Neurol. Sci. 2019; 396, 25–29. doi: 10.1016/j.jns.2018.10.024.

6. Castaldo I., De Rosa M., Romano A. et al. DNA damage signatures in peripheral blood cells as biomarkers in prodromal huntington disease. Ann. Neurol. 2019; 85, 296–301. doi: 10.1002/ana.25393.

7. Weiss A., Träger U., Wild E.J. et al. Mutant huntingtin fragmentation in immune cells tracks Huntington’s disease progression. J. Clin. Invest. 2012; 122, 3731–3736. doi: 10.1172/JCI64565.

8. Mastrokolias A., Ariyurek Y., Goeman J.J. et al. Huntington’s disease biomarker progression profile identified by transcriptome sequencing in peripheral blood. Eur. J. Hum. Genet. 2015; 23, 1349–1356. doi: 10.1038/ejhg.2014.281.

9. Przybyl L., Wozna-Wysocka M., Kozlowska E. & Fiszer A. What, When and How to Measure—Peripheral Biomarkers in Therapy of Huntington’s Disease. Int J Mol Sci 2021 Feb 4;22(4):1561. doi: 10.3390/ijms22041561.

10. Díez-Planelles C., Sánchez-Lozano P., Crespo M.C. et al. Circulating microRNAs in Huntington’s disease: Emerging mediators in metabolic impairment. Pharmacol. Res. 2016;108, 102–110. doi: 10.1016/j.phrs.2016.05.005.

11. Gaughwin P.M., Ciesla M., Lahiri N., Tabrizi, S.J., Brundin P., Björkqvistet M. Hsa-miR-34b is a plasma-stable microRNA that is elevated in pre-manifest Huntington’s disease. Human molecular genetics 2011; 20, 2225–2237. doi: 10.1093/hmg/ddr111.

12. Hoss A. G., Lagomarsino V. N., Frank S., Hadzi T. C., Myers R.H., Latourelle J.C. et al. Study of plasma-derived miRNAs mimic differences in Huntington’s disease brain. Mov. Disord. 2015;30, 1961–1964. doi: 10.1002/mds.26457.

13. Ferraldeschi M., Romano S., Giglio S. et al. Circulating hsa-miR-323b-3p in Huntington’s Disease: A Pilot Study. Front Neurol 2021; 12, 657973. doi: 10.3389/fneur.2021.657973.

14. Stepanov G. A., Filippova J.A., Komissarov A.B., Kuligina E.V., Richter V.A., Semenov D.V. Regulatory Role of Small Nucleolar RNAs in Human Diseases. BioMed Research International 2015, e206849. doi: 10.1155/2015/206849.

15. Cavaillé J., Hadjiolov A.A. & Bachellerie J. P. Processing of mammalian rRNA precursors at the 3’ end of 18S rRNA. Identification of cis-acting signals suggests the involvement of U13 small nucleolar RNA. Eur J Biochem 1996; 242, 206–213. doi: 10.1111/j.1432-1033.1996.0206r.x.

16. Kroh E. M., Parkin R. K., Mitchell P.S. & Tewari M. Analysis of circulating microRNA biomarkers in plasma and serum using quantitative reverse transcription-PCR (qRT-PCR). Methods 2010; 50, 298–301. doi: 10.1016/j.ymeth.2010.01.032.

17. Cawthon, R. M. Telomere measurement by quantitative PCR. Nucleic Acids Res. 2002; 30, e47. doi: 10.1093/nar/30.10.e47.

18. Goksuluk D., Korkmaz S., Zararsiz G. & Karaagaoglu A. E. EasyROC: An Interactive Web-tool for ROC Curve Analysis Using R Language Environment. The R Journal 2016; 8, 213–230.

19. Kang J., Tang Q., He J. et al. RNAInter v4.0: RNA interactome repository with redefined confidence scoring system and improved accessibility. Nucleic Acids Res 2022;50, D326–D332. doi: 10.1093/nar/gkab997.

20. Bouchard-Bourelle P., Desjardins-Henri C., Mathurin-St-Pierre D. et al. snoDB: an interactive database of human snoRNA sequences, abundance and interactions. Nucleic Acids Res 2020;48, D220–D225. doi: 10.1093/nar/gkz884.

21. Lang B., Armaos A. & Tartaglia G. G. RNAct: Protein–RNA interaction predictions for model organisms with supporting experimental data. Nucleic Acids Res 2019;47, D601–D606. doi: 10.1093/nar/gky967.

22. Szklarczyk D., Gable A. L., Lyon D. et al. STRING v11: protein-protein association networks with increased coverage, supporting functional discovery in genome-wide experimental datasets. Nucleic Acids Res 2019;47, D607–D613. doi: 10.1093/nar/gky1131.

23. Shannon P., Markiel A., Ozier O. et al. Cytoscape: a software environment for integrated models of biomolecular interaction networks. Genome Res 2003;13, 2498–2504. doi: 10.1101/gr.1239303.

24. Wickham H. Ggplot2: Elegant Graphics for Data Analysis. Springer-Verlag New York, 2016. ISBN 978-3-319-24277-4, https://ggplot2.tidyverse.org.

25. Genetic Modifiers of Huntington’s Disease (GeM-HD) Consortium. Electronic address: & Genetic Modifiers of Huntington’s Disease (GeM-HD) Consortium. CAG Repeat Not Polyglutamine Length Determines Timing of Huntington’s Disease Onset. Cell 2019;178, 887–900.e14. doi: doi: 10.1016/j.cell.2019.06.036.

26. Martí, E. RNA toxicity induced by expanded CAG repeats in Huntington’s disease. Brain Pathology 2016;26, 779–786. doi: 10.1111/bpa.12427.

27. Tsoi H., Lau T. C.-K., Tsang S.-Y. et al. CAG expansion induces nucleolar stress in polyglutamine diseases. Proc Natl Acad Sci U S A 2012;109, 13428–13433. doi: 10.1073/pnas.1204089109.

28. Nalavade R., Griesche N., Ryan D. P., Hildebrand S. & Krauss S. Mechanisms of RNA-induced toxicity in CAG repeat disorders. Cell Death Dis 2013;4, e752. doi: 10.1038/cddis.2013.276.

29. Tsoi H. & Chan H. Y. E. Roles of the nucleolus in the CAG RNA-mediated toxicity. Biochim Biophys Acta 2014;1842, 779–784. doi: 10.1016/j.bbadis.2013.11.015.

30. Kalita K., Makonchuk D., Gomes C., Zheng J.-J. & Hetman M. Inhibition of nucleolar transcription as a trigger for neuronal apoptosis. J Neurochem 2008;105, 2286–2299. doi: 10.1111/j.1471-4159.2008.05316.x.

31. Parlato R., Kreiner G., Erdmann G. et al. Activation of an Endogenous Suicide Response after Perturbation of rRNA Synthesis Leads to Neurodegeneration in Mice. J. Neurosci. 2008;28, 12759–12764. doi: 10.1523/JNEUROSCI.2439-08.2008.

32. Bordi I., Umeton R., Ricigliano V. A. G. et al. A mechanistic, stochastic model helps understand multiple sclerosis course and patho-genesis. Int J Genomics 2013, 910321. doi: 10.1155/2013/910321.

33. Adak D., Majumder A. & Bairagi N. Mathematical perspective of Covid-19 pandemic: Disease extinction criteria in deterministic and stochastic models. Chaos Solitons Fractals 2021; 142, 110381. doi: doi: 10.1016/j.chaos.2020.110381.

